# Actin and myosin dynamics during epithelial remodeling in avian gastrulation

**DOI:** 10.64898/2025.12.05.692587

**Authors:** Yu Ieda, Carole Phan, Olinda Alegria-Prévot, Aurélien Villedieu, Jérôme Gros

**Affiliations:** Institut Pasteur, Université de Paris, CNRS UMR3738, Developmental and Stem Cell Biology Department, F-75015 Paris, France; Collège Doctoral, Complexité du vivant, Sorbonne Université, Paris, France

**Author notes:** Contributed equally (Carole Phan, Olinda Alegria-Prévot).

## Abstract

Epithelial remodeling is powered by contractile forces exerted by the actomyosin cytoskeleton. In invertebrates, pulsatile contractile flows of the medio-apical actomyosin cortex have been shown to be critical in promoting junction contraction that ultimately drive cell rearrangements, apical constriction, and cell extrusion. However, how actomyosin dynamics drives epithelial remodeling in amniotes remains poorly understood. In this study, we generated transgenic quail lines reporting actin and myosin and investigate their dynamics in gastrulating embryos. We show that during this process, epithelial remodeling events are closely associated with the contraction of junctional myosin. Although we observe medio-apical contractile flows, those appear to contribute to junction contraction by promoting junctional myosin recruitment. Notably, by characterizing live and apoptotic cell extrusions and their associated actomyosin dynamics, we provide new insights into the cellular processes underlying primitive streak formation and the emergence of germ layers.

## Introduction

Embryonic development involves the incessant remodeling of cellular ensembles, arranging them into properly shaped tissues and organ primordia. Embryonic cells, initially organized as epithelial sheets, constantly balance adhesive forces that maintain epithelial cohesiveness with tensile forces that drive the shape changes underlying tissue morphogenesis. The actomyosin cytoskeleton plays a pivotal role in regulating this balance. On the one hand, fibrous actin (F-actin) stabilizes E-cadherin–based adhesive complexes at apical junctions by interacting with their intracellular domains. On the other hand, the same actin microfilaments serve as a scaffold for bipolar filaments of myosin-II molecular motors, whose processive activity enables filament sliding and thus generates contractile forces at the cortex. This tug-of-war at adhesive contacts creates cortical tension that drives local deformations (for review see^1^) and even large-scale tissue flows when actomyosin contraction is coordinated and propagated throughout tissues^2–4^.

Actomyosin dynamics and its pulsatile behavior have been shown to be critical for eliciting different morphogenetic events. Studies in Drosophila have shown that two pools of actin and myosin can be distinguished in epithelial cells: a junctional and a medio-apical population. Medial-to-junctional pulsatile actomyosin flows tend to pull and shrink junctions, eventually leading to polarized T1 neighbor exchange if flows are biased toward a specific junction^5,6^, and can even drive T2 cell extrusion if flows are persistent and directed toward all junctions^7,8^. Constriction of the apical area can also occur if flows are oriented toward the medial cortex, as shown during Drosophila ventral-furrow formation and dorsal closure; in this case, medial myosin contraction pulls on junctions through a ratchet-like process^9–11^. In apical constriction, pulsatile contractile flows drive the process differently from the “purse-string” model proposed for neurulating amphibian embryos, where the contraction of a junctional actomyosin belt would constrict the apical area^12–14^. Recently, studies analyzing apical constriction during chick and frog neural tube closure have identified pulsed contraction of the medio-apical cortex^15,16^. Furthermore, analyses of apical constriction in neuroectodermal cells and bottle cells during Xenopus gastrulation showed that the decrease in apical area correlates more strongly with an increase in medio-apical rather than junctional actomyosin, suggesting that medio-apical actomyosin accumulation may be the predominant driver of apical constriction in these cells^17,18^. Although *in vitro* studies have demonstrated that junctional myosin can generate tension and promote apical constriction^19–21^, whether apical constriction is driven more efficiently by circumferential belts or by medio-apical contraction remains unclear^22,23^ and likely depends on context. Actomyosin pulsed contraction has also been observed during the compaction of eight-cell preimplantation mouse embryos^24^. However, due to technical challenges associated with high-resolution dynamic imaging of live mouse embryos, characterizing actomyosin dynamics during complex morphogenesis in mammals, and amniotes in general, has remained a challenge. The Japanese quail (*Coturnix japonica*) has emerged as a particularly suitable amniote model for studying dynamic processes. Its small adult size and short generation period (8 weeks) are decisive advantages over the chicken model as a lab animal model and numerous transgenic quail lines, allowing to capture cellular and signaling dynamics^25–30^, have been generated. With its flat 2-dimensional geometry and ease of *ex ovo* culture at early stages of embryogenesis, the gastrulating quail embryo is an excellent system to study epithelial remodeling dynamics. During gastrulation, epithelial cells divide and rearrange throughout the epiblast^31^, while cells at the posterior margin of the embryonic domain engage in convergence-extension movements leading to the formation of the primitive streak^4,32,33^, where they eventually ingress to form the germ layers (mesoderm and endoderm). Here, we develop two transgenic quail lines reporting actin and myosin and analyze their dynamics during gastrulation. Although we identify medio-apical and pulsatile actomyosin behavior, we find that epithelial remodeling underlying convergent extension and cell ingression during germ-layer internalization at the primitive streak is strongly associated with the contraction of junctional myosin. Furthermore, we characterize the dynamics of apoptotic cell extrusions in the epiblast and show that their dynamics differ both qualitatively and quantitatively from that of ingressing cells at the primitive streak. Altogether, by identifying actomyosin dynamics during early quail embryogenesis, our study sheds new light on the cellular events underlying primitive streak formation and the emergence of germ layers.

## Results

### Generation of transgenic quail lines reporting actin and myosin

To visualize actin and myosin in live embryos, we generated two independent transgenic quail lines expressing, under the control of the human Ubiquitin-C (hUBC) promoter, the LifeAct peptide^27^ fused to a mNeonGreen fluorescent reporter (Tg (hUBC:LifeAct-mNeonGreen)) and the chicken myosin regulatory light chain *MYL9* fused to a tdTomato fluorescent reporter (Tg(hUBC:Myosin-tdTomato)), using a lentiviral approach^34^ (see Methods). Stable lines carrying one copy of each transgene were maintained as homozygous and interbred to obtain double-homozygous animals. Their embryos developed normally, exhibiting stable and high expression of the transgenes from stage Eyal-Giladi & Kochav (EGK) XI^35^ to at least Hamburger & Hamilton (HH) stage 15^36,37^ (Figure 1A**–**D). In these embryos, LifeAct-mNeonGreen and Myosin-tdTomato proteins were enriched at apical junctions in epithelial cells of the epiblast at stage EGK XI and stage HH 5, and of somites and neural tube at stage HH 15 (Figure 1A**–**D’). In stage EGK XI embryos, actin and myosin are strongly enriched at cell–cell junctions of epiblast cells and weakly distributed at the medio-apical cortex, without obvious organization (Figure 1B). Cells also exhibit short actin-rich apical protrusions (1 – 3 µm) reminiscent of microvilli (Figure 1B’). To confirm that these transgenic lines correctly report actin and myosin localization, we treated stage HH 1 embryos with H1152 and Latrunculin A, which inhibit myosin phosphorylation and actin polymerization, respectively. H1152 treatment abolished myosin phosphorylation, as revealed by immunofluorescence, and markedly reduced Myosin-tdTomato enrichment at cell junctions (Supplementary Figure 1A**–**B). Similarly, Latrunculin A treatment caused a drastic reduction of F-actin at epiblast cell junctions, as revealed by phalloidin staining and LifeAct-mNeonGreen imaging (Supplementary Figure 1C**–**D). Together, these results demonstrate that the Tg(hUBC:LifeAct-mNeonGreen) and Tg(hUBC:Myosin-tdTomato) quail lines faithfully report actin and myosin localization.

**Figure 1.**
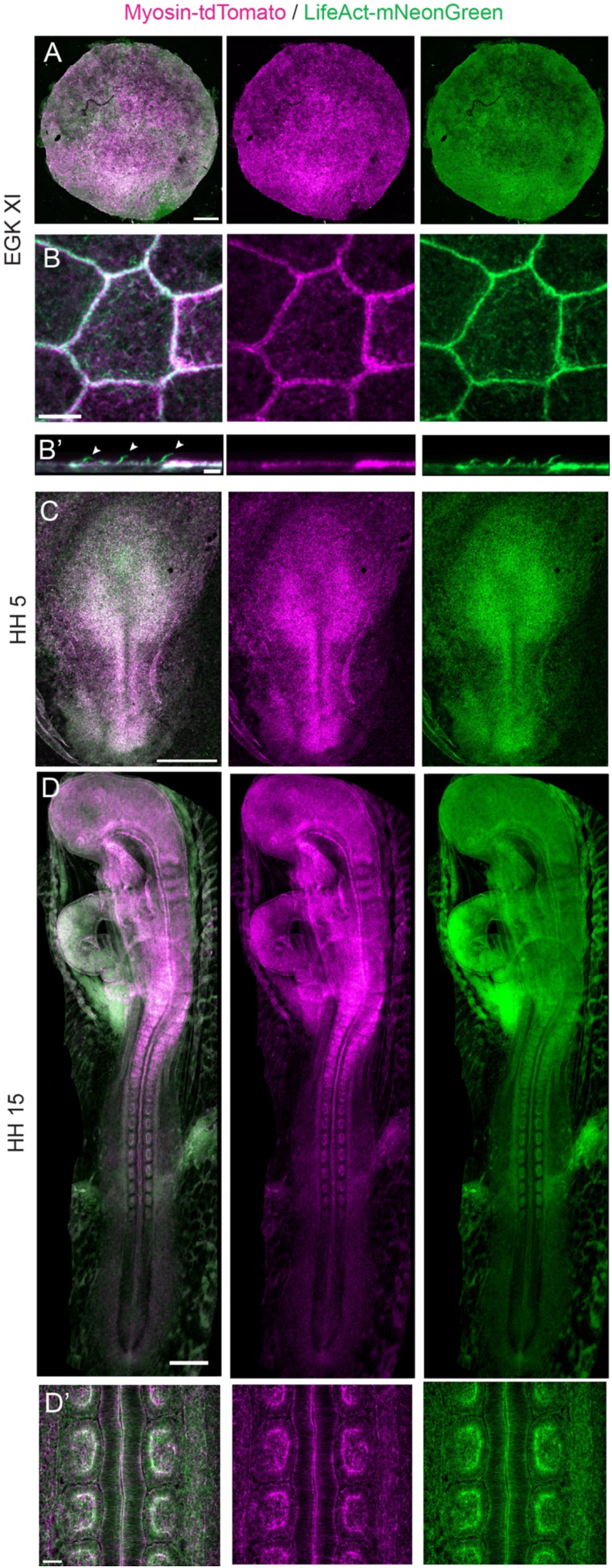
Transgenic quail lines reporting actin and myosin. (**A–D**) Transgenic embryos expressing Myosin-tdTomato (magenta) and LifeAct-mNeonGreen (green) at stages EGK XI (A, B), HH 5 (C), and HH 15 (D). (B–B’) Higher-magnification view of epithelial cells in the epiblast from (A) and corresponding orthogonal xz-section through the cell apical surface; white arrowheads mark microvilli (B’). (D’) Higher-magnification view of the neural tube and somites in the stage HH 15 embryo shown in (D). Scale bars: 500 µm (A, C, D), 50 µm (D’), 5 µm (B) and 2 µm (B’).

### Actin and Myosin dynamics during epithelial rearrangements

Next, we characterized actomyosin dynamics during gastrulation. To this end, we used the fast Airyscan 2 module of a Zeiss LSM 980, which enables super-resolved and rapid acquisition without detrimental bleaching. We first focused on epithelial cells in the center of the stage EGK XI epiblast, before the large-scale rotational tissue flows accompanying primitive streak formation (Figure 2A). In the absence of morphogenetic movements, we found that actomyosin contraction is pulsatile, with myosin and actin clusters appearing transiently at the very apical cortex (Figure 2B, and Movie S1 left column). These transient accumulations produced minor cell deformations that were not stabilized over time and did not lead to stable rearrangements. Thus, as in other systems, the actomyosin cortex of quail early epiblast cells exhibit pulsed contraction. As cells divided, myosin and actin accumulated at the cleavage furrow before its contraction (Figure 2C, and Movie S1 right column). After division, daughter cells intercalated between neighbors, as described previously^31^. These intercalations were associated with increased myosin intensity and subsequent junction contraction of neighboring junctions, suggesting that the T1 events driving daughter-cell intercalation may be active processes, potentially initiated by the dividing cell or the resulting daughter cells.

**Figure 2.**
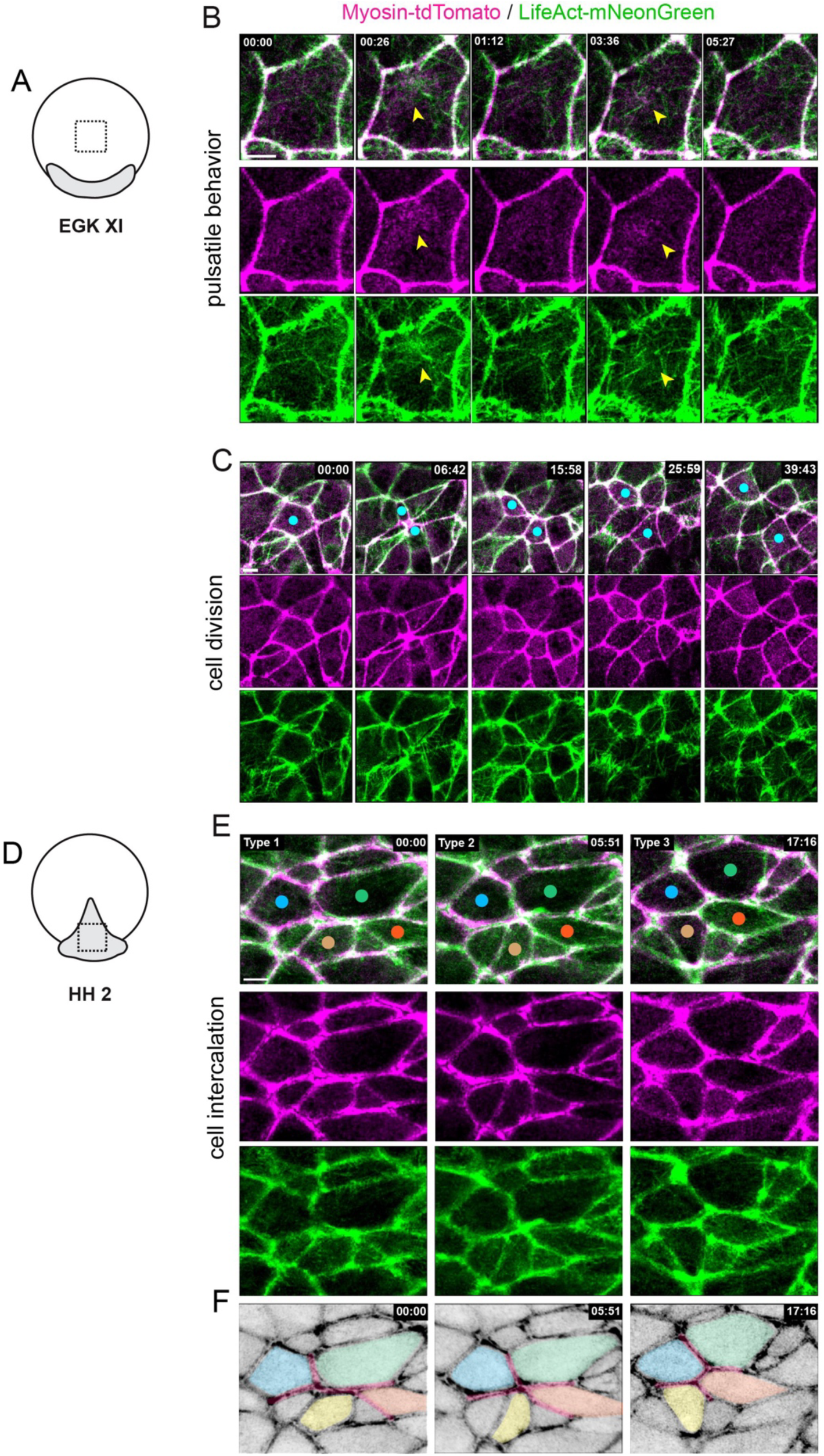
Actomyosin dynamics in epithelial rearrangement before and during avian gastrulation. (**A**) Schematic showing the region of live imaging (boxed) corresponding to the time series in (B) and (C). (**B–C**) Time series acquired in a central region of the epiblast of stage EGK XI transgenic embryos expressing Myosin-tdTomato (magenta) and LifeAct-mNeonGreen (green) showing myosin and actin pulsatile behavior in (B) and actin and myosin dynamics during cell division (C). Yellow arrowheads in (B) point at the transient enrichment of actin and myosin. Light blue dots in (C) point at a dividing cell and its daughters. (**D**) Sketch showing the location (squared region) of live-imaging acquisitions shown in (E). (**E–F**) Time series in the posterior margin of a stage HH 2 (E) showing actin and myosin dynamics during mediolateral intercalations underlying primitive streak elongation. Cells are tracked with colored dots. (F) Cell tracking displayed in the grayscale myosin channel from panel (E), with cell tracks shown in colors matching the overlaid dots. Junctions undergoing neighbor-exchange events are indicated by red overlays. Time is shown in min:sec. Scale bars: 10 µm (B**–**C, E).

We then examined actomyosin dynamics during the convergent extension movements that shape the primitive streak. Contraction of supracellular actomyosin cables at the posterior margin has been shown to drive cell mediolateral intercalation^4,33^, but their fine dynamics have not been characterized. We therefore live-imaged the posterior embryonic margin of stage HH 2 – 3 embryos. The reporters localized to cell junctions but were more enriched than in stage EGK XI epiblast: the junctional cortex was thicker, with actin-myosin fibers often distinguishable and aligned parallel to epithelial junctions. Notably, the individual cortex of two apposed cells could be readily distinguished using the Myosin-tdTomato fusion protein (Figure 2D–E). Myosin intensity was higher at anterior and posterior junctions, which predominantly contracted, promoting polarized T1 events (Figure 2E–F, and Movie S2 arrowheads). Higher spatiotemporal resolution imaging (every ∼5 s) revealed that over time myosin intensity increased at contracting junctions through the recruitment of actin and myosin (Movie S3 arrowheads). This recruitment was associated with rapid medio-apical contractile flows that emerged near the contracting junction and merged with it. Thus, we conclude that Myosin-tdTomato intensity is a good predictor of junction contraction, and that polarized recruitment of medio-apical actin and myosin at junctions underlies the convergent extension movements during primitive streak formation.

### Actin and myosin dynamics during cell ingression at the primitive streak

By stage HH 3, as epiblast cells converge to form the primitive streak, they ingress, generating the underlying mesenchymal mesendoderm through an epithelial-to-mesenchymal transition (EMT). Basement membrane destabilization has been reported as the first step initiating cell ingression in chicken embryos^38–40^. However, how cells subsequently disengage from apical adhesive contacts has not been characterized. While the apical ends of ingressing cells cannot be reliably tracked using a previously generated transgenic quails carrying a membrane-bound eGFP reporter (Tg(hUBC:memGFP))^30^ because of intense membrane blebbing (Movie S4), this becomes readily possible using the Myosin-tdTomato and LifeAct-mNeonGreen reporters. We found that cell ingression is characterized by the progressive formation of an inner myosin junctional belt, accompanied by concomitant apical constriction (Figure 3A**–**B’, and Movie S5 left column). As the junctional myosin belt contracted, cells became oblong and then circular until they extruded from the epiblast. Constrictions are asynchronous between cells of the forming primitive streak and often created gaps, which are followed by local cortex thickening in neighboring cells. Imaging embryos expressing both Myosin-tdTomato and membrane-bound eGFP revealed that blebbing coincides with the formation of these gaps, suggesting that they arise from cortex– membrane detachment caused by the rapid constriction (Movie S4). To quantify cell ingression dynamics, we segmented ingressing cells over time using the Myosin-tdTomato signal. We did not observe a ratchet-like mechanism in which constriction pauses and restarts, as described during Drosophila and mouse gastrulation^10,41^. Instead, cells constricted steadily at an average rate of 3.74 ± 1.73 µm².min⁻¹ for ∼20 min (Figure 3C**–**E). High-temporal-resolution imaging revealed that as observed during cell rearrangements, myosin recruitment to the contracting circumferential belt is associated with intense medio-apical contractile flows, during which myosin clusters appear near cell junctions, coalesce with actin fibers and eventually merge with junctional actin and myosin (Movie S6). Altogether, these analyses indicate that cells ingress through the formation of a circumferential actomyosin belt, whose dynamics are consistent with a purse-string closure mechanism that appears to be facilitated by medio-apical actomyosin activity.

**Figure 3.**
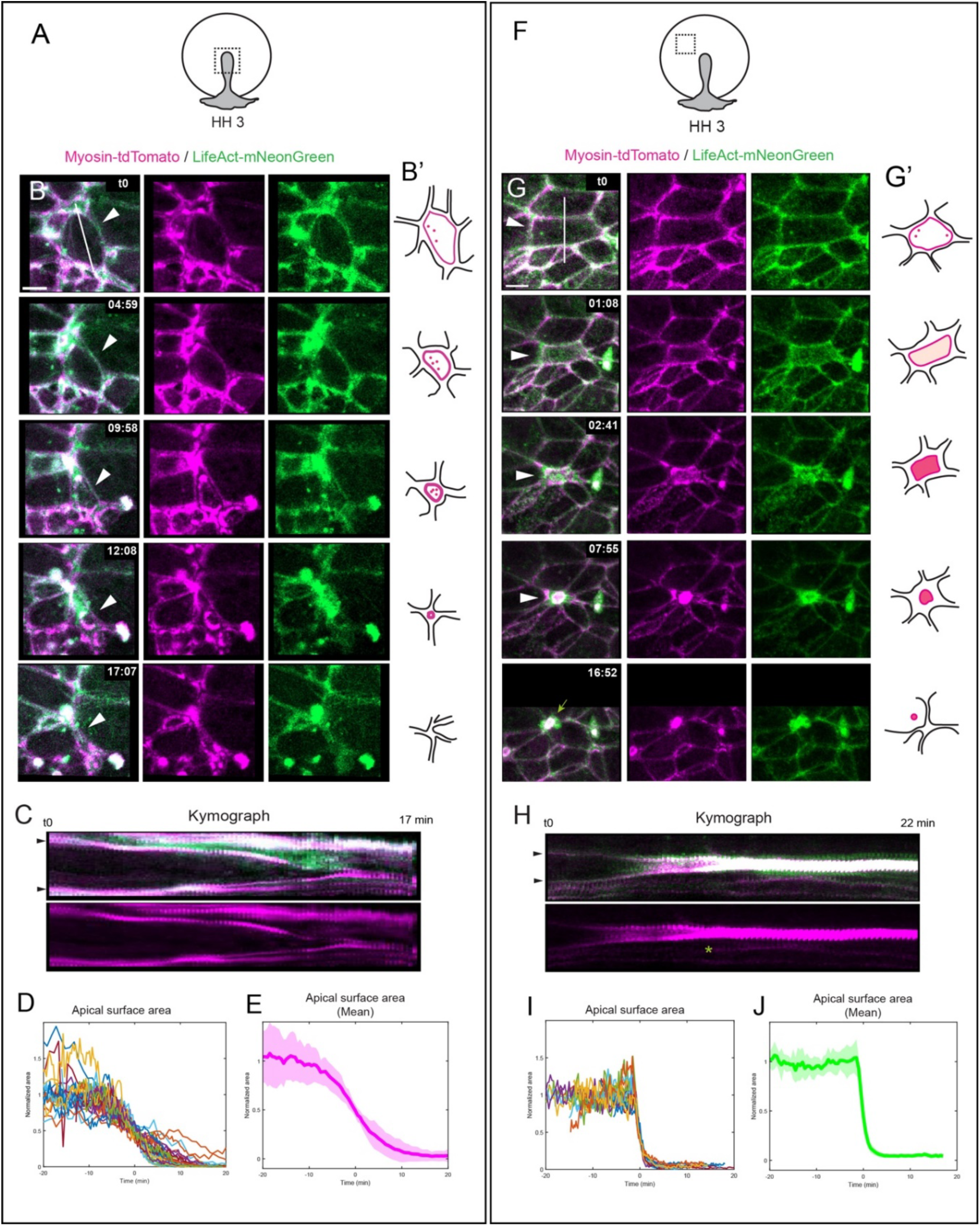
Actomyosin dynamics during cell ingression at the primitive streak and cell extrusion across the epiblast. (**A, F**) Schematic showing the region live imaged (boxed) shown in (B and G). (**B, G**) Time series acquired at the primitive streak (B) or in the central epiblast (G) of a stage HH 3 transgenic embryo expressing Myosin-tdTomato (magenta) and LifeAct-mNeonGreen (green), showing myosin and actin dynamics during cell ingression (B) and cell extrusion (G). White arrowheads indicate an ingressing cell (B) or an extruding cell. (**B’, G’**) Schematic depicting myosin distribution in the ingressing (B’) or extruding cell (G’). (**C, H**) Kymograph generated along the white line in (B, G). Black arrowheads point to cell boundaries. (**D– J**) Quantification of apical surface area of cell over time during ingression (D) and extrusion (J): normalized individual traces (D, I) and mean values (E, J) from n = 33 cells across 4 embryos (D, E) and n= 24 cells across 4 embryos (I, J). Curves are centered on each inflection point (see Methods). Error bands in (E, J) represent standard deviation. Time is shown in min:sec in (B, G). Scale bars: 5 µm.

Ingression of mesendodermal cells has been proposed to be initially scattered across the epiblast^42,43^ and amplified only at the primitive streak through a *NODAL*-dependent positive feedback loop^43^. To test whether we could visualize such early and scattered individual ingressions, we imaged large regions in the center of the epiblast away from the streak in Tg(hUBC:Myosin-tdTomato) embryos (Movie S7). In these movies, cells divided extensively, and we observed rare ingressions occurring prior to the onset of gastrulation movements, but not after they had begun (Movie S8, and Supplementary Figure 2A). These early ingressions proceed much more slowly than those at the forming primitive streak, with durations ranging from 56 min to 5 h 39 min (n = 23 cells across 2 embryos; Movie S8 and Supplementary Figure 2A). To further investigate the fate of these cells, we electroporated the anterior epiblast stage EGK XI memGFP transgenic embryos with a cytoplasmic tdTomato reporter and examined tdTomato⁺ cells 12 hours later, before the emergence of mesendodermal cells from the primitive streak.

In these embryos, tdTomato^+^ cells could be observed underneath the epiblast (Supplementary Figure 2B**–**B’). These cells were either rounded or associated with the epithelializing hypoblast as revealed by immunofluorescence for the tight junction protein: ZO-1 and the transcription factor protein: FOXA2/HNF3𝛽, which are expressed by the hypoblast at this stage (Supplementary Figure 2C)^44^. These results suggest that early ingressing cells might not be related to gastrulation but to polyingression. This process, during which the hypoblast cells and primordial germ cells (PGCs) are generated, begins during intrauterine development and ends just before germ layer formation begins^35,45–48^. Consistent with this, we find that 24 hours after electroporation, cytoplasmic eGFP^+^ cells originating from the anterior epiblast could be observed in the germinal crescent and expressed CVH (chicken vasa-homolog), which is specifically expressed by PGCs (Supplementary Figure 2D**–**E)^49^. Altogether these results show that early cell ingressions across the epiblast produce precursors of the hypoblast and PGCs rather than mesendoderm cells and are therefore related to polyingression and not to gastrulation.

### Actin and myosin dynamics during apoptotic cell extrusion in the epiblast

Large-scale movies of the epiblast in Tg(hUBC:Myosin-tdTomato) embryos (Movie S7) revealed numerous cell extrusions as gastrulation movements are taking place. However, these events differed both qualitatively and quantitatively from those observed at the primitive streak. High-resolution imaging of Tg(hUBC:Myosin-tdTomato; hUBC:LifeAct-mNeonGreen) embryos showed that these extrusions are characterized by the sudden accumulation of actin and myosin across the entire apical surface in the form of a plaque, which abruptly contracted, disengaging all neighbor contacts (Figure 3F**–**H, and Movie S5 right column). This was followed by a localized increase in actin and myosin in the surrounding cells. This contrasts with the progressive contraction of a myosin junctional belt observed in ingressing cells at the forming primitive streak. Segmentation revealed that this extrusion mode is significantly faster than that of ingressing cells at the primitive streak, with an average constriction rate of 21.0 ± 12.2 µm².min⁻¹ and a duration of ∼5 min (compare Figure 3E and Figure 3J, see Supplementary Figure 5A**–**B). As cells extruded, the apical actomyosin plaque was expelled apically and drifted in the space between the epiblast and the vitelline membrane, suggesting cell fragmentation (Figure 3G, and Supplementary Figure 3C**–**D). To follow the fate of these extruding cells, we electroporated Tg(hUBC:Myosin-tdTomato) embryos with cytoplasmic eGFP and performed 3D time-lapse imaging. Analyzing movies along the Z axis revealed that ingressing cells electroporated in the primitive streak region, display a high basal protrusive activity and then ingress basally once apical constriction is complete, consistent with basement membrane destabilization preceding apical disengagement^38–40^ (Supplementary Figure 3A**–**B, Movie S9 left column, and Movie S10). In contrast, cells electroporated away from the primitive streak fragmented upon apical actomyosin accumulation, with eGFP^+^ debris expelled both apically and basally (Supplementary Figure 3C**–**D, Movie S9 right column, and Movie S11). To confirm that these extrusions are apoptotic, we performed immunofluorescence for cleaved Caspase-3 (CASP3) on Tg(hUBC:Myosin-tdTomato; hUBC:LifeAct-mNeonGreen) embryos (Figure 4A**–**B). We observed CASP3^+^ cells scattered across the epiblast, confirming widespread apoptotic events across the epiblast during gastrulation (Figure 4A). Notably, cells exhibiting apical Myosin-II plaques were strongly cleaved CASP3^+^ (Figure 4B) and a myosin cable extending basally could be observed, similar to what has been described during apoptosis in neural tube closure^50^ (Figure 4C white arrows, and Movie S11). Treatment with Camptothecin, a topoisomerase-1 inhibitor that generates double strand breaks and induces apoptosis^51,52^, drastically increased the number of cleaved CASP3^+^ cells compared to control embryos (Figure 4D**–**E, G**–**H). Live imaging revealed an increase in epithelial extrusions whose dynamics were comparable to those observed across the epiblast (20.0 ± 10.0 µm²·min⁻¹ over ∼5 minutes; Figure 4M–N, Supplementary Figure 5C–D, and Movie S12). Conversely, treating embryos with Q-VD-OPh, a potent pan-caspase inhibitor which efficiently inhibits apoptosis in avian embryos^4,50^, markedly reduced the number of cleaved CASP3-positive cells compared to control DMSO- and Camptothecin-treated embryos. It also abolished virtually all extrusions in the anterior epiblast (Figure 4D**–**H, Supplementary Figure 4A**–**C, and Movie S13 top). Notably, cell ingression at the primitive streak was unaffected both qualitatively and quantitatively: ingression still proceeded via formation of a junctional myosin belt, and the average constriction rates in control (4.38 ± 2.10 µm²·min⁻¹) and Q-VD-OPh– treated embryos (3.40 ± 2.20 µm²·min⁻¹) did not differ significantly (Supplementary Figure 5E–J, and Movie S13 below). Altogether these results demonstrate that these fast extrusions are apoptotic events and that apical actomyosin contraction occurs downstream of caspase activation.

**Figure 4.**
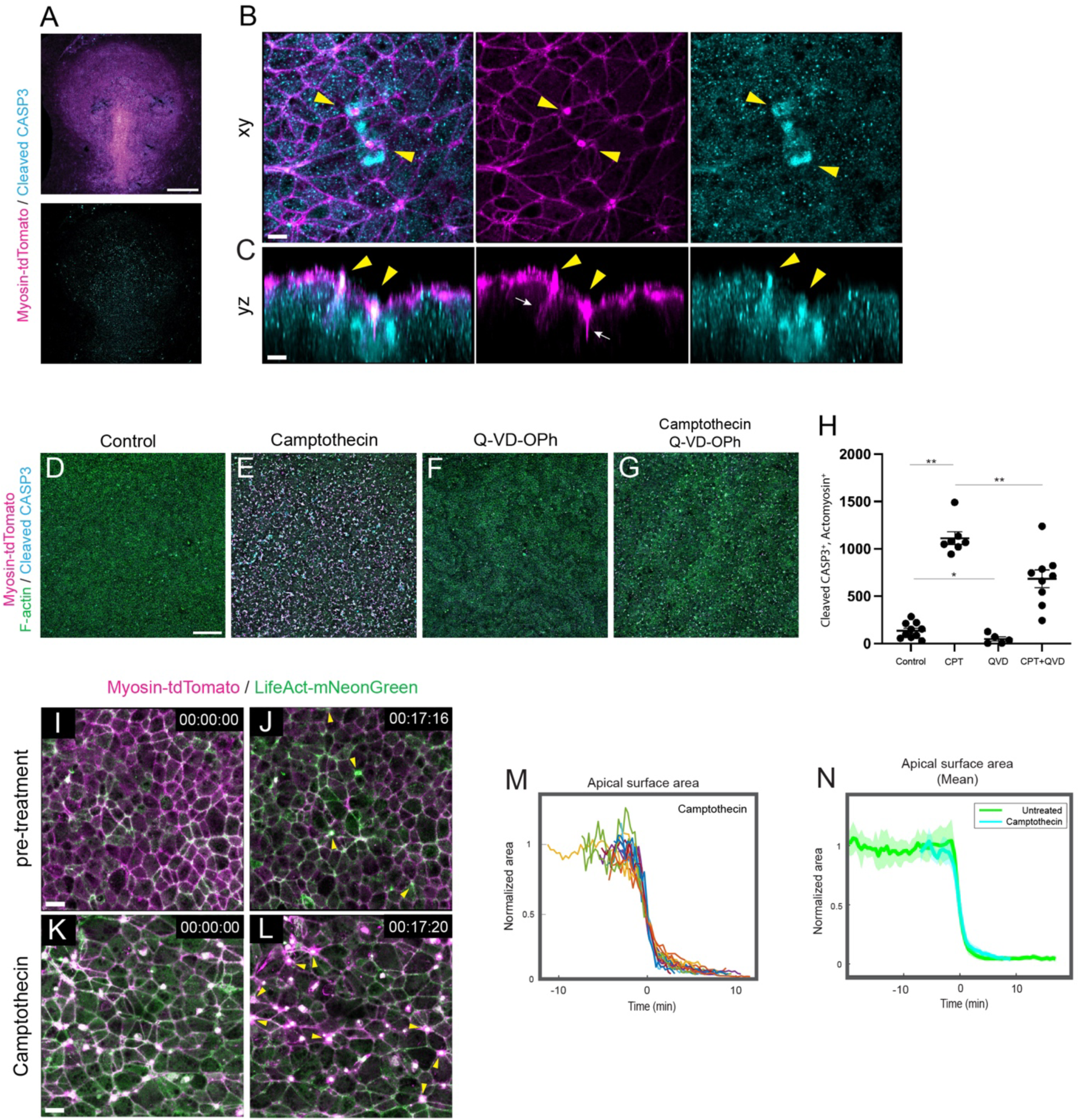
Scattered cell extrusions during gastrulation are apoptotic events downstream of caspases. (**A**) Immunofluorescence for cleaved Caspase-3 (cleaved CASP3, cyan) in a stage HH 3 Myosin-tdTomato transgenic embryo (Myosin-tdTomato in magenta). (**B**) Higher magnification of the central epiblast region from (A), showing co-localization of apical myosin plaques and cleaved CASP3 (yellow arrowheads). (**C**) Orthogonal yz-slice of the image in (B), highlighting extruding cells. Yellow arrows indicate apical myosin plaques; white arrows denote apicobasal myosin cables. (**D–G**) Stage HH 3 Myosin-tdTomato transgenic embryos treated for 5 hours with DMSO (D), Camptothecin (E), Q-VD-OPh (F), or Q-VD-OPh and Campthotecin (G) followed by staining with phalloidin and cleaved CASP3 antibody. (**H**) Quantification of Actomyosin⁺; cleaved CASP3⁺ puncta density (counts / mm^2^) in central epiblast regions after drug treatment (see Methods). Control (DMSO), n = 10 embryos; Camptothecin (CPT), n = 7; Q-VD-OPh (QVD), n = 6; Q-VD-OPh + Camptothecin (CPT+QVD), n = 9. \*\**p* < 0.01 (Control vs. Camptothecin); \*\**p* < 0.01 (Camptothecin vs. Camptothecin + Q-VD-OPh); \**p* < 0.05 (Control vs. Q-VD-OPh). One-way ANOVA. (**I–L**) Time series of a central epiblast region from a stage HH 3 transgenic embryo expressing Myosin-tdTomato (magenta) and LifeAct-mNeonGreen (green), captured before (I**–**J) and after Camptothecin treatment (K–L). Yellow arrowheads mark extruding cells. (**M–N**) Quantification of apical surface area over time during cell extrusion after Camptothecin treatment showing normalized individual cell curves (M) and averaged curves (N, and see Supplementary Figure 5C) plotted together with quantifications from untreated embryos (see Figure 3J), from 16 cells from 3 embryos. Curves are aligned on each cell’s inflection point (see Methods). Error bands in (N) represent standard deviation. Time is shown in hour:min:sec in (I**–**L). Scale bars: 500 µm (A), 100 µm (D), 10 µm (I, K) and 5 µm (B**–**C).

## Discussion

In this study, we generated transgenic quail lines enabling the capture of actin and myosin dynamics in live embryos, at high spatiotemporal resolution. Using these transgenic lines, we characterize actomyosin dynamics during key epithelial remodeling events, including cell division, polarized intercalations, live and apoptotic cell extrusions, underlying avian gastrulation (Figure 5A**–**C). Although we observed cell extrusions away from the primitive streak, their dynamics differ markedly. Our results indicate that these extrusions are associated either with hypoblast/PGC formation at early, pre-gastrulation stages or with apoptotic events during gastrulation and are therefore unrelated to mesendoderm formation (Figure 5D).

**Figure 5.**
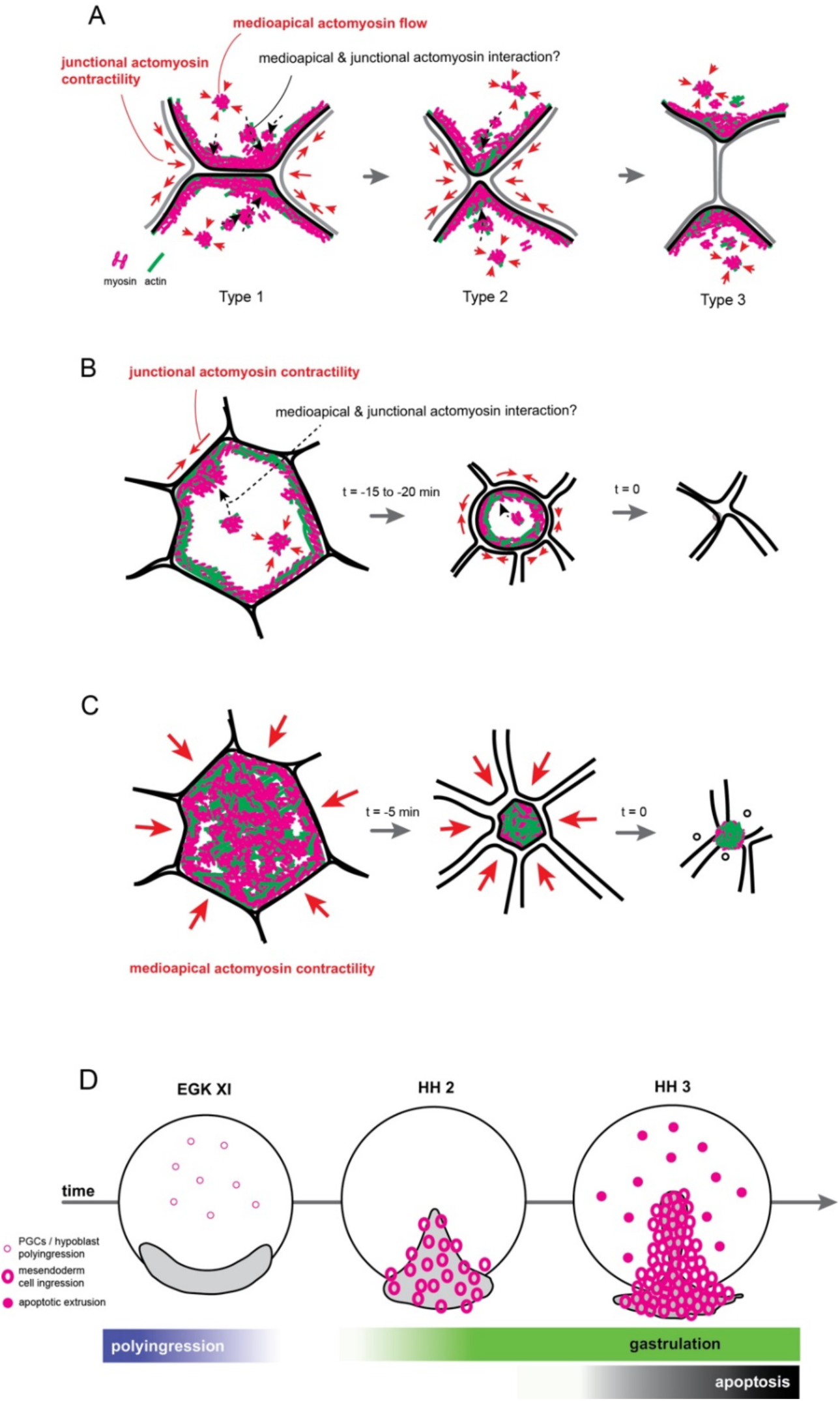
Schematic model of actomyosin behaviors during epithelial remodeling dynamics in avian gastrulation. (**A–C**) Schematic representation of actin and myosin dynamics during epithelial remodeling in avian gastrulation. (**A**) Junctional apical actomyosin and medio-apical actomyosin flows contribute to junctional contractility that drives cell intercalation during primitive streak formation. Time progresses from left (Type 1) to right (Type 3), illustrating a T1 transition. (**B**) Junctional apical actomyosin, together with medio-apical actomyosin flows, appears to drive cell ingression. Time progresses from left to right. (**C**) Caspase-dependent, strong accumulation of medio-apical actomyosin drives apoptotic cell extrusion from the avian epiblast, leaving a residual apical plaque at the tissue surface. Time progresses from left to right. Red arrows depict contractile forces; gray arrows indicate time progression; dashed black arrows denote molecular recruitment. Myosin molecules are shown in magenta and F-actin in green. (**D**) Different types of cell extrusion during early avian development. Shallow magenta rings indicate polyingression events associated with primordial germ cell (PGC) formation and hypoblast formation prior to gastrulation. Thick magenta rings represent mesendoderm cell ingression through the primitive streak during gastrulation. Thick magenta dots denote apoptotic cell extrusion occurring throughout the embryonic territory during gastrulation.

Recently a transgenic quail line expressing a LifeAct-EGFP transgene has been reported^27^. Here, we report a similar transgenic line expressing a LifeAct-mNeonGreen in combination with a new Myosin-tdTomato transgenic line. Comparing the localization of these two transgenes in the same embryos, we find that while LifeAct-mNeonGreen is expressed widely at the cortex, the Myosin-tdTomato reporter provides unique insights into contractile behaviors, with its intensity being a good predictor of cortical contraction, as illustrated during cell division, polarized cell rearrangements, live and apoptotic cell extrusions. Furthermore, we find that these epithelial remodeling events largely associate with junctional myosin contraction. High-resolution spatiotemporal imaging reveals intense medio-apical actomyosin activity. These movies suggest that medio-apical contractile flows contribute to junction contraction by merging with junctional myosin but also potentially by exerting medial pulling forces on the junctions that might facilitate their contraction (Figure 5A**–**B). However, quantifying the specific contributions of junctional and medial actomyosin pools will be needed to assess their respective roles in epithelial remodeling events. The identification that medio-apical contractile flows regulate **–**or are associated with**–** apical constriction in various developmental contexts and models challenged the initial proposition that apical constriction is mediated by a purse-string mechanism. In the case of cell ingression at the primitive streak, we observe the formation of a clear actomyosin circumferential belt whose contraction dynamics are consistent with a purse-string mechanism: apical constriction occurs steadily, as the junctional myosin belt reduces its circumference until cells extrude from the epiblast (Figure 5B). This observation contrasts with recent findings in the murine primitive streak, where apical constriction during cell ingression has been associated with a pulsed ratchet-like behavior^41^. It should be noted, however, that in that study, actin and myosin dynamics were not characterized, and apical constriction was quantified using a ZO-1-GFP protein fusion reporter. Notably, in mouse, apical constriction was described to last 25 – 90 min, whereas in quail, we find that it lasts ∼20 min, suggesting that ingression in mice and birds might indeed involve different subcellular mechanisms.

The ingression of mesendodermal cells has been proposed to occur scattered across the epiblast and amplified only at the primitive streak^42,43^. However, the fate of these early ingressing cells had not been characterized. Although we identify rare ingressions before the onset of the large-scale rotational tissue flows that accompany primitive streak formation, we find that these ingressions are slower (occurring over several hours) than those at the primitive streak. Furthermore, by following the fate of these cells using the electroporation of an eGFP reporter, we find that they express FOXA2, which is expressed by the anterior hypoblast at this stage^44^, and indeed associate with it . Moreover, we show that some of these electroporated cells can also be found in the germinal crescent and express CVH, which is specifically expressed by PGCs. The hypoblast and PGCs segregate from the epiblast through the ventral shedding of cells in a process named “polyingression”^48^. Based on transmitted light and electron microscopy studies^35,45–48^, polyingression has been described to initiate during intrauterine development and to proceed progressively from posterior to anterior, persisting anteriorly during the first 6 – 8 hours of post-oviposition development. Our dynamic imaging, which shows that ingressions in the anterior epiblast can no longer be observed after ∼8 hours of development, together with our lineage analysis, thus support that these early ingression events are “polyingressions” of hypoblast and PGC precursors rather than the scattered ingression of mesendodermal cells related gastrulation^43^. Once the primitive streak has emerged, we did find extruding cells scattered across the epiblast, but we show that they are apoptotic events which proceed much faster than ingressions observed at the primitive streak. Furthermore, we show that these extrusions are not associated with the formation of a circumferential actomyosin belt but instead involve an apical plaque of actomyosin that abruptly contracts, provoking the fragmentation of the cell in a caspase-dependent manner (Figure 5C). Our results are therefore consistent with other reports showing that actomyosin-driven epithelial extrusion during apoptosis is downstream of caspase activation^53–56^.

## Methods

### Avian embryos

All experimental methods, animal husbandry and transgenesis for transgenic quails were performed in accordance with the guidelines of the European Union 2010/63/UE, approved by the Institut Pasteur ethics committee authorization: #dha210003, and under the GMO agreement: #2432. Fertilized Japanese quail eggs were ordered from commercial sources (Chanteloup, France), and transgenic Japanese quail eggs were produced in the lab. Quail fertilized eggs were kept at 15 °C to store.

### *Ex ovo* culture and *ex ovo* time-lapse photography of quail embryos

For microsurgeries and live cell imaging, embryos were incubated for the desired stage at 38.5 °C. Before each manipulation, the embryo was carefully immersed drop by drop in Hank’s Balanced Salt Solution (HBSS). Japanese quail eggs were incubated at 38.5 °C up to the appropriate developmental stage. For the very early stage (stage EGK XI), the eggs were equilibrated at room temperature for 1 hour. All embryos were staged according to the Hamburger and Hamilton or the Eyal-Giladi & Kochav classification system^35–37^. Embryos were then collected (stage EGK XI) using a paper filter ring and cultured on a semi-solid nutritive medium made of thin chicken albumen, agarose (0.2%), glucose and NaCl, as described previously with its epiblast surface and vitelline membrane facing the semi-solid nutritive medium^30,57^. They were inverted and placed on a 50 mm uncoated glass-bottom dish (MatTek, #P50-G-0-30F) or a 6-well uncoated glass-bottom dish (MatTeck, #P06-G-1.5-20-F) covered by the same fresh semi-solid medium for live-cell imaging at 38.5 °C using an inverted confocal microscope (either of Zeiss LSM 700 or LSM 900 or of LSM 980 Airyscan 2) with X5, X10, X20 or X40 objectives in the temperature-controlled chamber at 38.5 °C.

### Generation of Myosin-tdTomato and LifeAct-mNeonGreen transgenic quail lines

Two transgenic lines were generated in this study (hUBC:Myosin-tdTomato and hUBC:LifeAct-mNeonGreen) by following previously published method^26^. Briefly, non-incubated Japanese quail eggs (*Coturnix japonica*) were windowed and a solution of high titer lentivirus was injected into the subgerminal cavity of stage EGK X embryos. Eggs were sealed with a plastic piece and paraffin wax. Injected eggs were incubated 38.5 °C, 56 % humidity until hatching. For the hUBC:Myosin-tdTomato line, a total of 386 embryos were injected with the lentivirus solution (titer 6.0 x 10^10^ /ml). Twenty-nine F0 mosaic founder successfully hatched and reached sexual maturity. They were bred to WT female and all three produced transgenic offspring (transmission rate: 6.7 %). One line was selected on the basis of a single copy of the transgene, checked by Southern Blot, and high intensity of the tdTomato signal. For the hUBC:LifeAct-mNeonGreen line, a total of 193 embryos were injected with lentivirus (titer 5.0 x 10^9^ /ml). Seventeen F0 mosaic founder successfully hatched and reached sexual maturity. All five produced transgenic offspring (transmission rate: 3.1 %) and one line was selected by Southern blot analysis for single transgene integration and high intensity of the mNeonGreen fluorescence. MemGFP-expressing transgenic quail (hUBC:membrane-bound eGFP) was maintained in the laboratory as previously described^30^. Myosin-tdTomato transgenic quail was subsequently crossed with the memGFP or LifeAct-mNeonGreen line to produce offspring carrying both transgenes.

### *Ex ovo* electroporation, drug treatments and live-cell imaging

*Ex ovo* electroporation was performed as previously described^30^. It introduced the cytosolic eGFP or tdTomato expressing plasmid (respectively pT2K-CAGGS-eGFP, or pT2K-CAGGS-tdTomato; 1.0 µg/µl in PBS(-)) to a stage EGK X to XII quail epiblast in a custom-made electroporation chamber using the SuperElectroporator NEPA21 type II® (NEPAGENE) electroporator and the electrode (NEPAGENE, #CUY700P3L or #CUY700P2L) with two poring pulses of 15.0 V for 8.00 ms, 50.0 ms delay, 10% decay rate and three transfer pulses of 5.00 V for 50.0 ms, 500 ms delay, 40% decay rate. Electroporated embryos were cultured by a modified version of EC culture system^57^ on either a plastic Petri dish with semisolid albumin/agarose (0.2 %) culture medium to keep them developing in a petri dish, or a 50 mm uncoated glass-bottom dish (MatTek, #P50-G-0-30F) or a 6-well uncoated glass-bottom dish (MatTeck, #P06-G-1.5-20-F) with same medium for live-cell imaging at 38.5 °C with humidity.

The albumen-based culture medium contains drugs in case of treatment: (S)-(+)- Camptothecin (Sigma-Aldrich, #C9911; 100 µM), Q-VD-OPh (Merck, #SML0063; 50 - 500 µM), H1152 (Tocris, 35 µM) and Latrunculin A (Sigma-Aldrich, #L5163; 50 µM), DMSO (Sigma-Aldrich, #D2650; 0.5 – 1.0 %). They were placed under either of Zeiss LSM 700 or LSM 900 or of LSM 980 Airyscan 2) with X2.5, X5, X10, X20 objectives for live-cell imaging acquisition inside the temperature-controlled chamber at 38.5 °C attached by a microscope device.

### Immunocytochemistry

For antibody staining in whole-mount embryos, samples were washed in PBS (-) and fixed in 4 % paraformaldehyde / PBS (-) at 4 °C for overnight. The fixed samples were rigorously washed in PBST (0.2 % bovine serum albumen, 0.02 % SDS, and 0.1 % Triton-X100 in PBS (-)). They were incubated with primary antibodies (see, Antibody list) in PBST or Can Get Signal™ immunostain Solution B (TOYOBO, #NKB-601) at room temperature for 2 hours or 4 °C for overnight. After rigorous washing, secondary antibody treatment was performed at 4 °C for overnight or at room temperature for 1 hours with antibodies in PBST or Can Get Signal™ immunostain Solution B. The samples are treated in Hoechst and phalloidin (BIOTIUM) in PBST. Embryos were then mounted, cover-slipped, and imaged under an inverted confocal microscope Zeiss LSM 900 or LSM 980 Airyscan 2.

### Actomyosin^+^; cleaved CASP3^+^ puncta density quantification

Myosin-tdTomato; LifeAct-mNeonGreen transgenic quail embryos (stage HH 3) were subjected to 5-hour drug treatments (DMSO 0.5%, Q-VD-OPh 50 µM, Camptothecin 100 µM, or Q-VD-OPh 50 µM + Camptothecin 100 µM) at 38.5 °C. Following treatment, the central epiblast (imaged region: 638.9 µm × 638.9 µm, excluding the primitive streak) was acquired from the dorso-apical side using a Zeiss LSM 900 microscope (X20 objective, zoom factor 0.5) after staining with anti-cleaved CASP3 and phalloidin (to enhance F-actin signal). Images were projected based on the apical surface detected from the phalloidin channel. After projection, each channel (Myosin-tdTomato, phalloidin, and cleaved CASP3) was binarized, merged, and used to detect overlapping signals corresponding to apical debris. Projection, signal detection, and debris quantification were automated and performed using MATLAB.

### Apical surface area quantification

After live-cell imaging acquisition, images were processed using maximum-intensity projection. Cells undergoing cell ingression or apoptotic extrusion were then manually tracked. Changes in apical area were quantified by manual frame-by-frame segmentation, using the Myosin-tdTomato signal to define cell junctions. For each curve of cell area over time, we fit a sigmoid curve *f(t)* that tends to zero when *t* tends to the infinite: 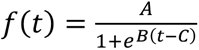

By derivating this function, we can extract the maximal apical constriction rate *S* at the inflection point:

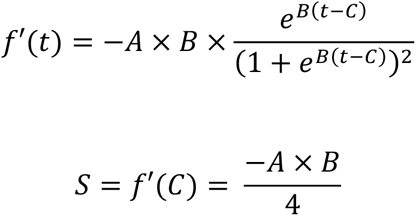

Normalized area was calculated by averaging area by *A*, which corresponds to the initial value of the fitted sigmoid. Each curve was aligned in time so that the time when the fitted sigmoid reaches its inflection point (corresponding to *C*) is set to 0. Maximum projection, cell tracking, and manual segmentation are done in Fiji. Rendering graphs with plots is performed in MATLAB.

### Antibody list

Rabbit Phosphorylated non-muscle myosin-II regulatory light chain 2 (Ser19/Thr18);

(1:250; Cell Signaling, #3674)

Mouse ZO-1 (1:250; Invitrogen, #33-9100);

Rabbit FoxA2/HNF-3β (1:500; Proteintech, #22474-1-AP);

Rat Chicken Vasa-homolog (CVH) (1:500; kind gift from Dr. Bertrand Pain’s group);

Rabbit Human/Mouse Active Caspase-3 (1:1000; R&D, #AF835);

Phalloidin 488 (invitrogen, #A12379).

## Acknowledgements

We thank Arthur Michaut, Alexander Chamolly, and Carolina Parada-Borja for valuable discussions and advice on experimental design and data analysis. Work in J.G.’s laboratory is supported by the European Research Council (ERC) under the European Union’s Horizon 2020 research and innovation programme (grant agreement no. 866186 to J.G.), the Agence Nationale de la Recherche (LabEx Revive), the Centre National de la Recherche Scientifique (CNRS), and the Institut Pasteur. Y.I. was supported by a stipend from the Pasteur–Paris University (PPU) International Doctoral Program, affiliated with Sorbonne University (École Doctorale Complexité du Vivant, ED515), and by a fourth-year PhD fellowship (Fin de thèse) from the Fondation pour la Recherche Médicale (FRM). For the purpose of open access, the authors have applied a CC BY public copyright license to any Author Accepted Manuscript version arising from this submission

## Author Contributions

The project was conceptualized by Y.I. and J.G. Y.I. and J.G. designed the experiments, and Y.I. performed the majority of them. Y.I. and J.G. analyzed the data and wrote the manuscript, including figure preparation. O.A.-P. and C.P. assisted with and conducted part of the experimental work. O.A.-P. and C.P. generated the Myosin:tdTomato; LifeAct-mNeonGreen transgenic quail line and managed the supply of transgenic eggs. A.V. and J.G. designed the quantitative image analysis strategies, statistical approaches, and data-analysis framework. A.V. developed the quantitative image-analysis pipeline, performed data quantification and visualization, and implemented the statistical analyses.

