## Supplementary Information for "Actin and myosin dynamics during epithelial remodeling in avian gastrulation"

- Supplementary Figures 1-5
- Supplementary Movie legends (Movies S1-S13)

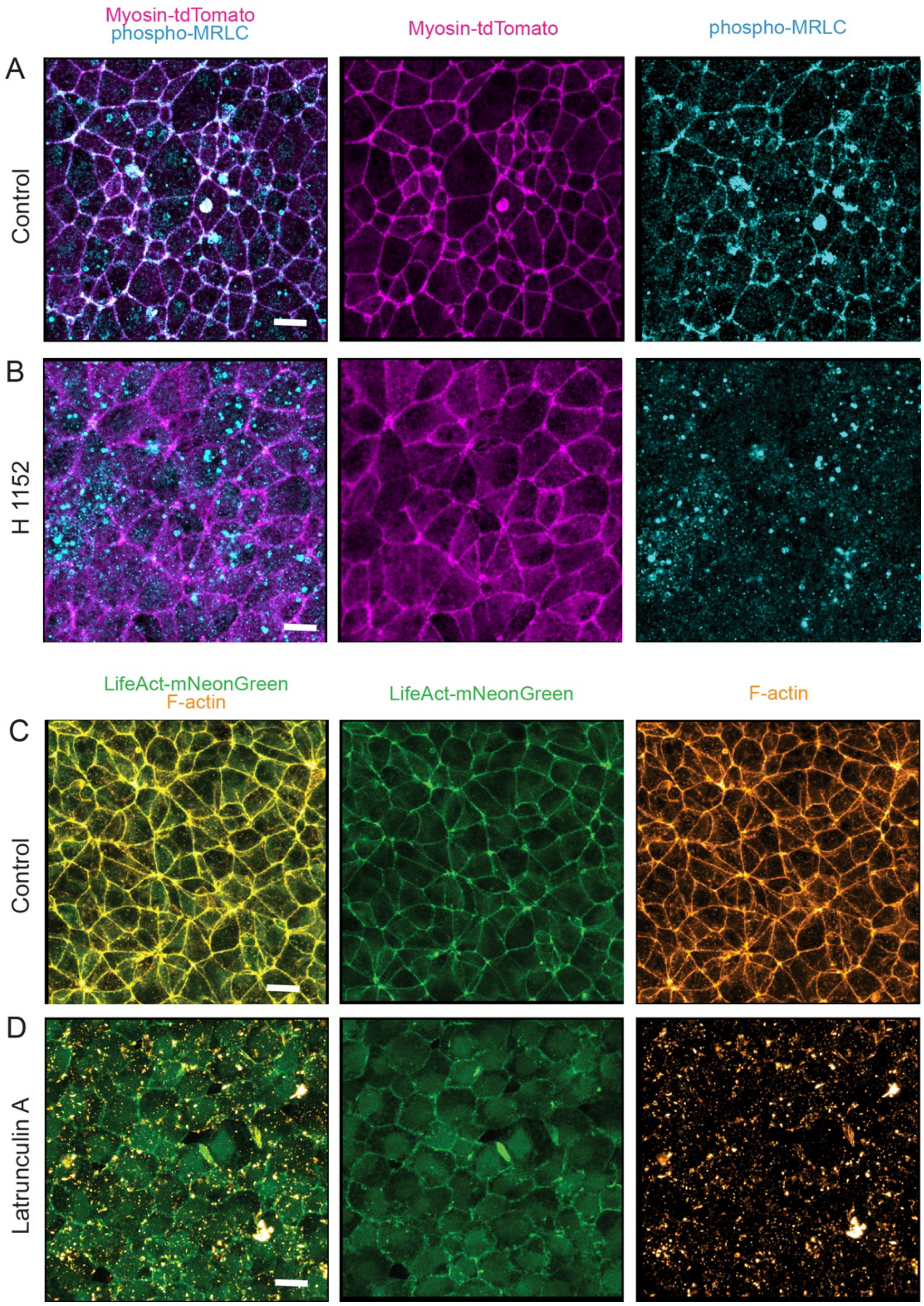


**Supplementary Figure 1. Myosin-tdTomato and LifeAct-mNeonGreen distribution upon pharmacological treatment.**
(**A–D**) Stage HH 1 transgenic embryos expressing Myosin-tdTomato (A–B) and LifeAct-mNeonGreen (C–D) treated with water (A), H1152 50 µM (B), DMSO 0.5% (C), and Latrunculin A 50 µM (D), and subjected to immunofluorescence for phosphorylated myosin-regulatory-light-chain-II (A–B; phospho-MRLC, cyan; Myosin-tdTomato, magenta; n = 3 embryos) and phalloidin staining for F-actin detection (C–D; F-actin, orange hot; LifeAct-mNeonGreen, green; n = 3 embryos).

Scale bars: 10 µm

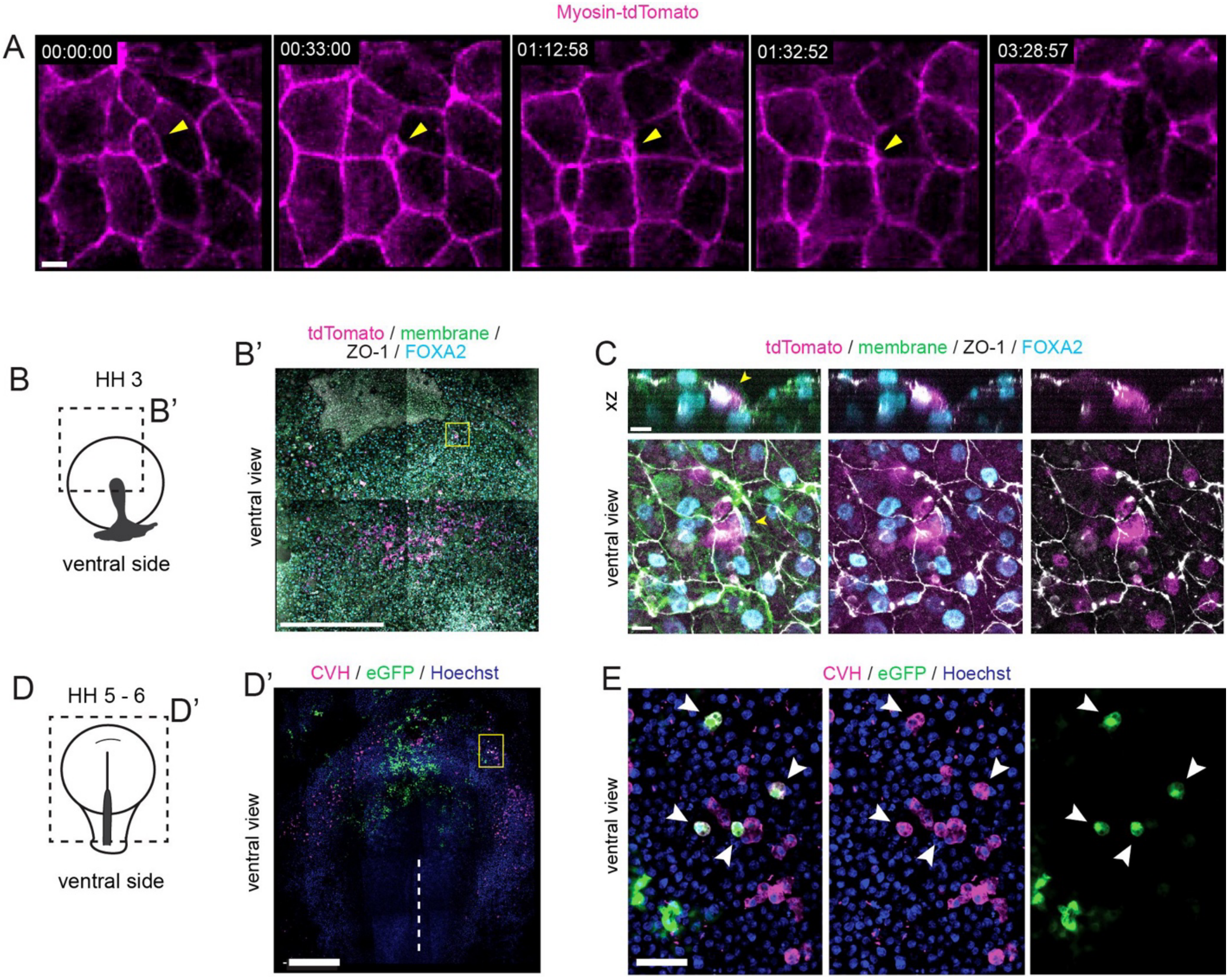


**Supplementary Figure 2. Dynamics and fate of early cell ingressing cells in stage EGK XI embryos.**
**(A)** Time series acquired in the central epiblast of a stage EGK XI transgenic embryo expressing Myosin-tdTomato (magenta), showing myosin dynamics during early cell ingression. Yellow arrowhead marks an early ingressing cell. **(B–C)** Stage HH 3 transgenic embryo expressing memGFP, electroporated at stage EGK XI in the anterior epiblast with a tdTomato transgene, and immunolabeled for ZO-1 (gray) and FOXA2 (cyan) after 12 hours (B–B’). (C) Higher magnification of the yellow-boxed region in (B’) and corresponding orthogonal xz-section. Yellow arrowheads indicate tdTomato$^+$/FOXA2$^+$ cells integrated within the ZO-1$^+$ epithelialized hypoblast. tdTomato$^+$ cells were observed in the hypoblast in 3 out of 4 embryos. **(D–E)** Stage HH 6 embryo electroporated in the anterior epiblast with an eGFP reporter at stage EGK XI, immunolabeled for CVH (magenta) and stained with Hoechst (blue) after 24 hours (D–D’). (E) Higher magnification of the yellow-boxed region in (D’). eGFP$^+$/CVH$^+$ cells were observed in 3 out of 3 embryos. Time is shown in hour:min:sec in (A).

Scale bars: 10 µm (A, C), 500 µm (B’, D’) and 50 µm (E).

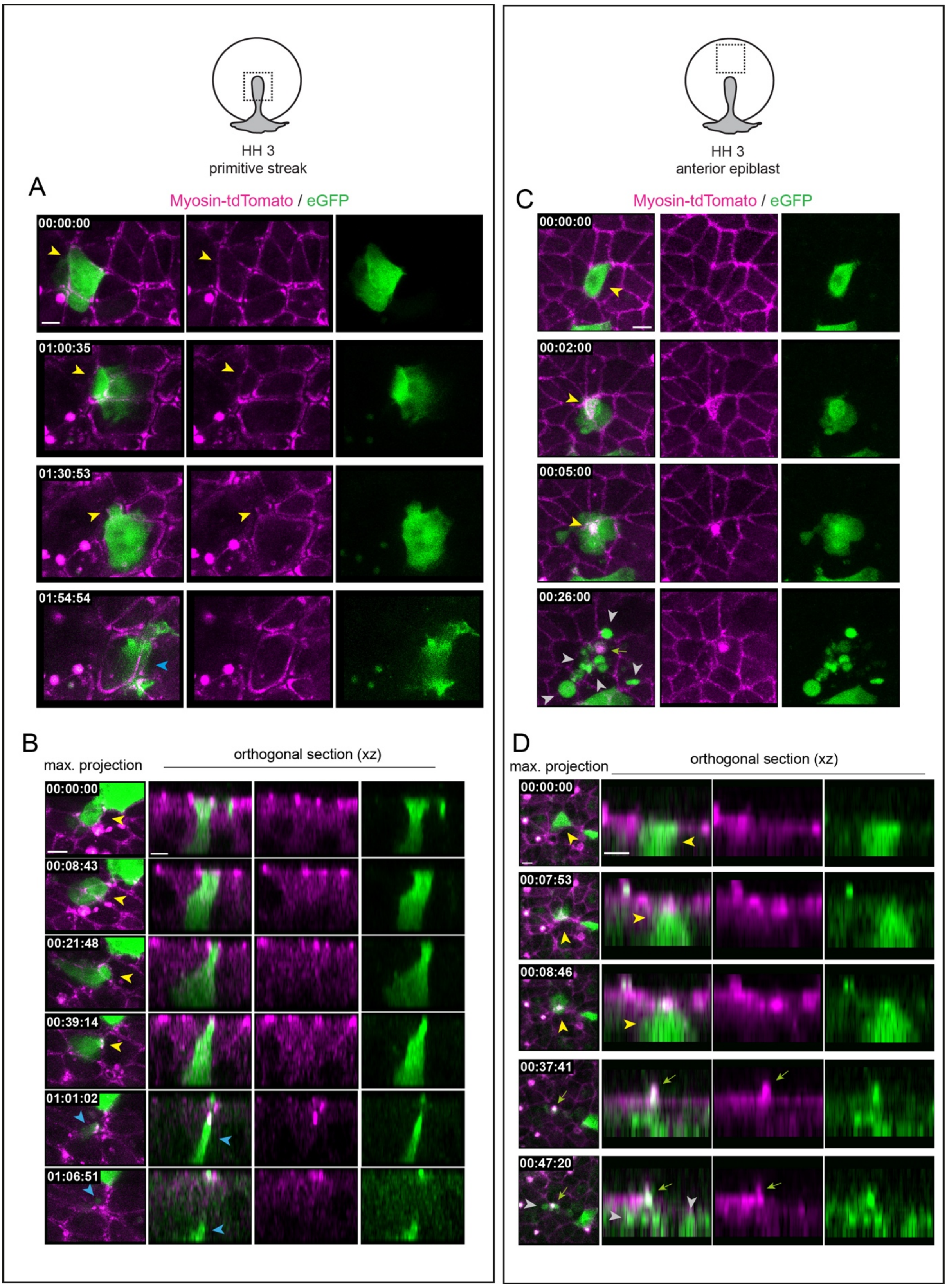


**Supplementary Figure 3. Cell shape changes during ingression at the primitive streak and extrusions across the epiblast.**
**(A)** Time series acquired at the primitive streak of a stage HH 3 transgenic embryo expressing Myosin-tdTomato (magenta) and electroporated with an eGFP reporter (green), showing cell shape changes during ingression. Yellow arrowheads indicate an eGFP$^{+}$ ingressing cell; the blue arrowhead marks the same cell after basal ingression. **(B)** Additional example of cell ingression at the primitive

streak, shown from an apical view and corresponding orthogonal xz-section. **(C)** Time series acquired in the central epiblast of a stage HH 3 transgenic embryo expressing Myosin-tdTomato (magenta) and electroporated with an eGFP reporter (green), showing cell shape changes during extrusion. Yellow arrowheads highlight the extruding cell; gray arrowheads indicate post-apoptotic fractures; green arrows mark apical myosin debris. **(D)** Additional example of apoptotic extrusion shown in apical view and corresponding xz optical cross-section. Yellow arrowheads highlight the extruding cell; gray arrowheads indicate post-apoptotic fractures; green arrows mark apical myosin debris. Time is shown in hours:min:sec in (A–D).

Scale bars: 5 μm.

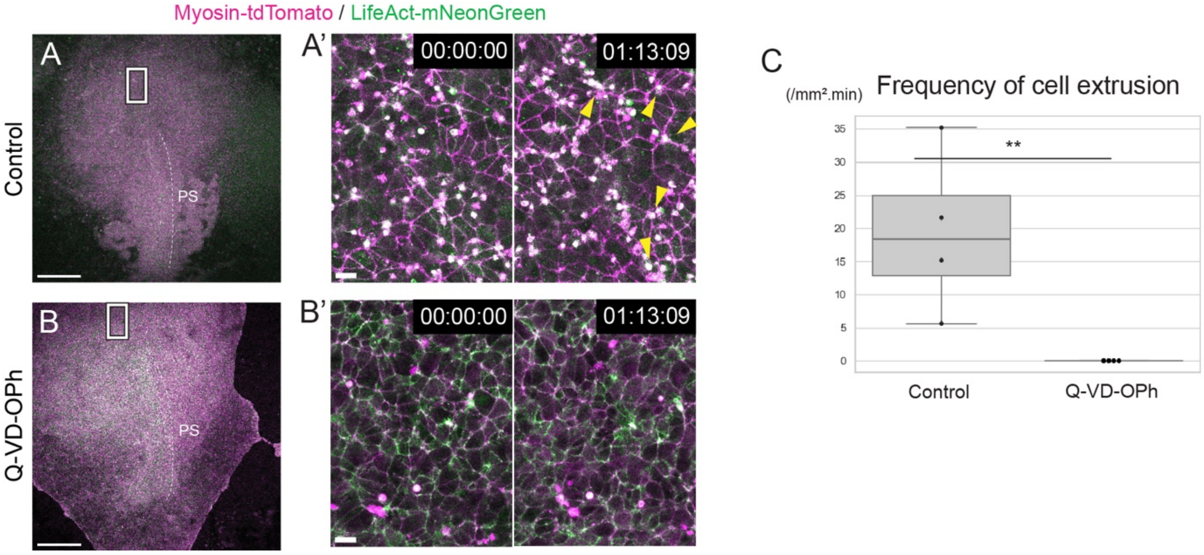


**Supplementary Figure 4. Caspase inhibition abolishes scattered cell extrusion in the epiblast of gastrulating embryos.**
(**A–B**) Time series of stage HH3 transgenic embryos expressing Myosin-tdTomato (magenta) and LifeAct-mNeonGreen (green), treated with DMSO (A) or Q-VD-OPh (B). White dashed lines indicate the primitive streak (PS). (**A'**, **B'**) Higher-magnification views of the regions boxed in (A–B). Yellow arrowheads mark apoptotic extrusions. Marged images are shown. **(C)** Frequency of cell extrusion in the epiblast (events/μm²·min) quantified by live imaging. Control: n = 4 embryos; Q-VD-OPh: n = 4 embryos. ***p* < 0.01, Mann–Whitney *U* test. Time is displayed as hour:min:sec in (A'**,** B').

Scale bar: 500μm (A**–**B) and 10 μm (A'**,** B').

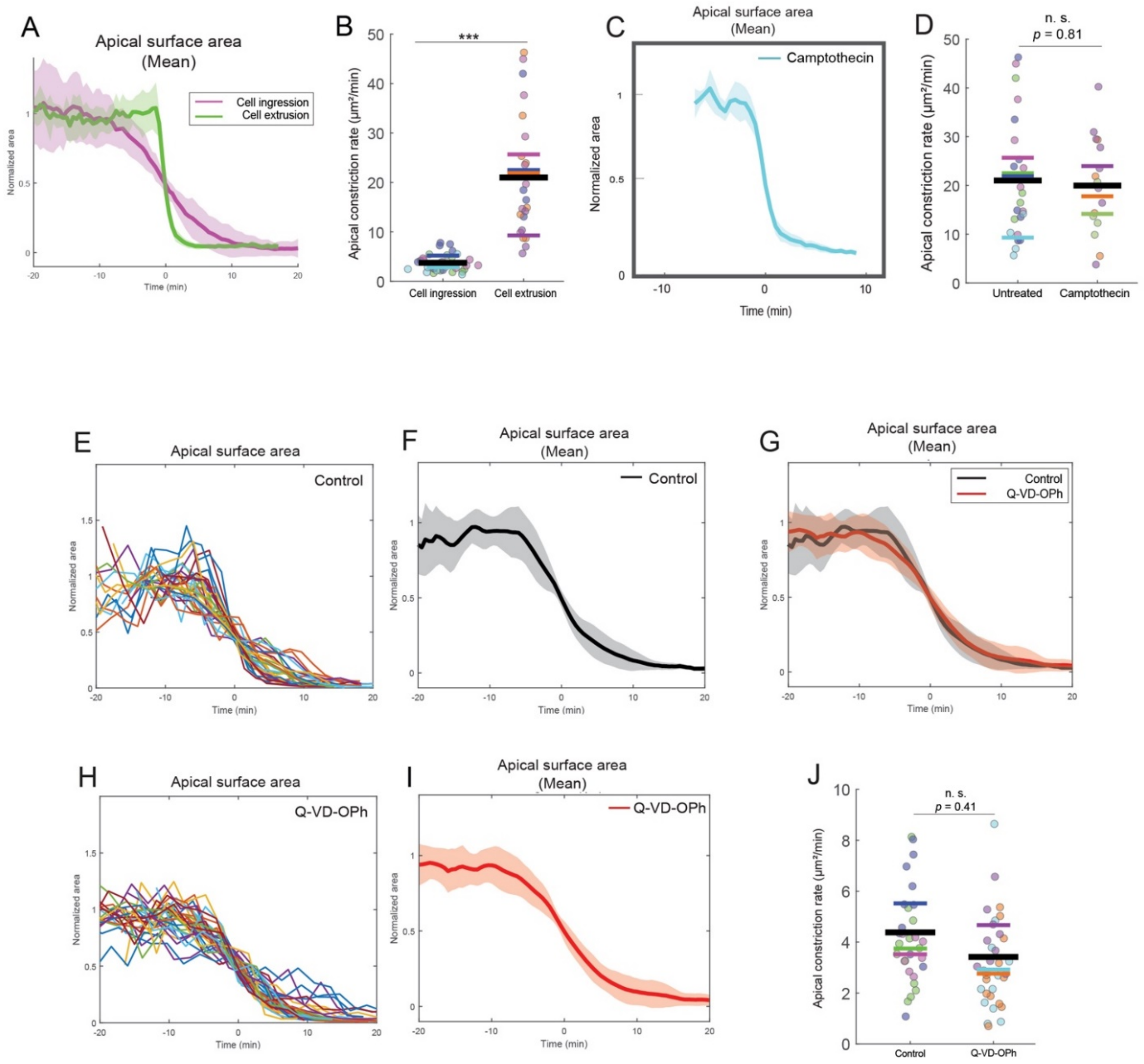


**Supplementary Figure 5. Supplementary quantifications and statistical analyses**

**(A–B)** Comparison of cell ingression and apoptosis surface area dynamics over time. (A) Comparison of the mean apical surface area curves for cell ingression (see Figure 3E) and apoptotic cell extrusion (see Figure 3J). (B) Scatter plots of apical constriction rates ($\mu m^2 \cdot min^{-1}$) during apical constriction associated with cell ingression (left) and cell extrusion (right). Each dot represents a single cell, and each color corresponds to a different embryo. Colored bars indicate the mean rate for each embryo, and the black bar indicates the overall mean for each condition. ***$p < 0.0001$ (linear mixed-effects model). (**C**)Mean apical surface area curve over time for cells undergoing extrusion after Camptothecin treatment (see Figure 4M). (**D**) Scatter plot of rate of apical constriction rate ($\mu m^2.min^{-1}$) during cell extrusion in untreated embryos (left) and in treated with Camptothecin. Each dot represents a single cell; colors correspond to different embryos; colored bars indicate embryo specific means and the black bar indicates the overall mean for each condition. $p = 0.808$ (Nested ANOVA); non-significance (n.s.) (**E–I**) Quantification of apical surface area over time during cell ingression in DMSO- (E–F) and Q-VD-OPh-treated embryos (H–I). Normalized individual cell curves **(**E, H**)** and corresponding mean curves (F, **I)** from 31 cells across 3 embryos for Control DMSO and from 35 cells across 3 Q-VD-OPh-treated embryos. Curves are aligned on each cell's inflection point (see Methods**).** Error bands in (F, I) represent standard deviation. (G) Comparison of mean curves between cell ingression from DMSO-treated embryos (F) and Q-VD-OPh-treated embryos (**I**). (**J**) Scatter plot of rate of apical constriction ($\mu m^2.min^{-1}$) during cell ingression in Control embryos (left) and in embryos after Q-VD-OPh treatment (right). Each dot represents an individual cell; colors indicate different embryos; colored bar indicates overall means for each embryo. Black bar indicates the overall mean for each condition. $p = 0.411$ (Nested ANOVA); non-significance (n.s.).

### Supplementary Movie Legends

**Movie S1. Pulsatile behavior and dynamics of actomyosin during mitosis at stage EGK XI**
Time-lapse imaging of a stage EGK XI transgenic quail epiblast expressing Myosin-tdTomato (magenta) and LifeAct-mNeonGreen (green). Left column: Pulsatile actomyosin behavior; white arrowheads mark cells of interest. Right column: Cell division (white arrowhead). From top to bottom: merged image, myosin channel, and actin channel.

Scale bar: 5 µm. Timestamp: hour:min:sec. Acquisition rate: 6.55 sec/frame.

**Movie S2. Actomyosin dynamics during polarized intercalations at the primitive streak**.
Time-lapse imaging of a stage HH 2 transgenic embryo expressing Myosin-tdTomato (magenta) and LifeAct-mNeonGreen (green) during polarized intercalations underlying primitive streak formation. White arrowheads at the start of the movie indicate myosin-enriched, contracting junctions. From top to bottom: merged image, grayscale myosin channel with overlaid dots from manual cell tracking, myosin channel, and actin channel.

Scale bar: 10 µm. Timestamp: hour:min:sec. Acquisition rate: 18.6 sec/frame.

**Movie S3. High spatiotemporal resolution movie of junction contraction during polarized intercalations at the primitive streak.**
Time-lapse imaging of a stage HH 3 transgenic embryo expressing Myosin-tdTomato (magenta) and LifeAct-mNeonGreen (green) during polarized intercalations underlying primitive streak formation. White arrowhead highlights contractile junctional cortices with showing medio-apical actomyosin recruitment. From top to bottom: merged image, myosin channel, and actin channel.

Scale bar: 5 µm. Timestamp: hour:min:sec. Acquisition rate: 4.6 sec/frame.

**Movie S4. Cell ingression at the primitive streak in memGFP; Myosin-tdTomato in stage HH 3 transgenic embryos.** Time-lapse imaging of a stage HH 3 transgenic embryo expressing Myosin-tdTomato (magenta) and memGFP (green) during cell ingression at the primitive streak. White arrowheads point to constricting cells; yellow arrowheads indicate the associated membrane blebbing. From left to right: merged image, myosin channel, and GFP channel.

Scale bar: 5 µm. Timestamp: hour:min:sec. Acquisition rate: 3.8 sec/frame.

**Movie S5. Distinct mechanisms of apical constriction in cell ingression and cell extrusion.**
Time-lapse imaging of a stage HH 3 transgenic embryo expressing Myosin-tdTomato (magenta) LifeAct-mNeonGreen (green) during cell ingression at the primitive streak (left column) and apoptotic extrusion in the epiblast (right column). From top to bottom: merged image, myosin channel, and actin channel. White arrowheads indicate the ingressing or extruding cell at movie onset.

Scale bars: 5 µm. Timestamp: hour:min:sec. Acquisition rate: 13.0 sec/frame (Cell ingression); 18.3 sec/frame (Cell extrusion).

**Movie S6. High spatiotemporal resolution movie of apical constriction during cell ingression at the primitive streak via junctional contraction and medio-apical activity.**
Time-lapse imaging of a stage HH 3 transgenic quail embryo expressing Myosin-tdTomato

(magenta) and LifeAct-mNeonGreen (green) during cell ingression. White arrowheads indicate cells of interest. White arrows indicate medio-apical actomyosin contractile flows.

Scale bar: 2 µm. Timestamp: hour:min:sec. Acquisition rate: 4.6 sec/frame.

**Movie S7. Apical myosin dynamics in the central epiblast from stage EGK XI to HH 3.**
Large-scale time-lapse imaging of a transgenic quail epiblast expressing Myosin-tdTomato (magenta) to visualize early ingression events from stage EGK XI to HH 3. White double-headed arrow: anterior–posterior axis at the onset of the movie (A, anterior; P, posterior). White bar: primitive streak at the end of the movie.

Scale bar: 100 µm. Timestamp: hour:min:sec. Acquisition rate: 3 min 19 sec/frame.

**Movie S8. Early cell ingression in anterior epiblast prior to gastrulation.**
Isolated early ingression event from Movie S7. Cell ingression prior to mesendoderm formation (stage EGK XI – HH 1). Left: Myosin-tdTomato (magenta). Right: manual tracking (yellow dot–line overlay on the grayscale myosin channel). Yellow arrowhead marks the ingressing cell.

Scale bar: 10 µm. Timestamp: hour:min:sec. Acquisition rate: 3 min 19 sec/frame.
Movie onset (00:00:00) corresponds to the beginning (00:00:00) of Movie S7.

**Movie S9. Cell shape changes during cell ingression at the primitive streak and cell extrusion across the epiblast.**
Time-lapse imaging of a stage HH 3 transgenic embryo expressing Myosin-tdTomato (magenta) electroporated with eGFP reporter (green) highlighting the cell shape changes during cell ingression at the primitive streak and cell extrusion across the epiblast. Left column: Cell ingression. Right column: Cell extrusion. White arrowheads indicate cells at movie onset. From top to bottom: merged image, myosin channel, and GFP channel.

Scale bar: 5 µm. Timestamp: hour:min:sec. Acquisition rate: 1 min/frame.

**Movie S10. Cell shape changes along the apicobasal axis during cell ingression at the primitive streak.**
Time-lapse imaging of a stage HH 3 transgenic embryo expressing Myosin-tdTomato (magenta) electroporated with eGFP reporter (green) highlighting the shape changes along the apicobasal axis during cell ingression at the primitive streak. White arrowhead marks an ingressing cell at movie onset. From top to bottom: merged maximum-projection image from the dorsal side; merged xz optical slice; merged xz optical slice of the myosin channel; and merged xz optical slice of the GFP channel.

Scale bar: 5 µm. Timestamp: hour:min:sec. Acquisition rate: 1 min/frame.

**Movie S11. Cell shape changes along the apicobasal axis during cell extrusion in the epiblast.**
Time-lapse imaging of a stage HH 4 transgenic embryo expressing Myosin-tdTomato (magenta) electroporated with eGFP reporter (green) highlighting the shape changes along the cell apicobasal axis during cell extrusion. White arrowheads indicate an extruding cell at movie onset. From top to bottom: merged maximum-projection image from the dorsal side; merged xz optical slice; merged xz optical slice of the myosin channel; and merged xz optical slice of the GFP channel.

Scale bar: 5 µm. Timestamp: hour:min:sec. Acquisition rate: 1 min/frame.

**Movie S12. Camptothecin-induced apoptotic cell extrusions in the epiblast**.
Time-lapse imaging of the central epiblast of a stage HH 3 transgenic embryo expressing Myosin-tdTomato (magenta) and LifeAct-mNeonGreen (green) before (left) and after Campthotecin treatment (right). Merged images are shown. Yellow arrowheads indicate extruding cells.

Scale bar: 10 μm. Timestamp: hour:min:sec. Acquisition rate: 14.8 sec/frame (Before treatment); 20.3 sec/frame (After Camptothecin treatment).

**Movie S13. Apototic cell extrusion and cell ingression at the primitive streak upon Q-VD-OPh treatment.**
Inhibition of apoptotic cell extrusion in the epiblast and unaffected cell ingression at the primitive streak upon Q-VD-OPh treatment, in a stage HH 3. Left column (DMSO control): anterior epiblast (top) and primitive streak (bottom). Right column (Q-VD-OPh): anterior epiblast (top) and primitive streak (bottom). Each column shows images from a single representative embryo (n = 4 embryos per condition). Yellow arrowheads indicate individual extrusion events. Merged channels: Myosin-tdTomato (magenta) and LifeAct-mNeonGreen (green).

Scale bar: 10 μm. Timestamp: hour:min:sec. Acquisition rate: 2 minutes 22 sec/frame.
